# Simulation of Gilbert Theory for Self-Association in Sedimentation Velocity Experiments: A Guide to Evaluate Best Fitting Models

**DOI:** 10.1101/2022.10.20.513061

**Authors:** GR Bishop, JJ Correia

## Abstract

There is a long tradition in the Biophysics community of using simulations as a means to understand macromolecular behavior in various physicochemical methods. This allows a rigorous means to interpret observations in terms of fundamental principles, including chemical equilibrium, reaction kinetics, transport processes and thermodynamics. Here we simulate data for the Gilbert Theory for self-association, a fundamental analytical ultracentrifuge (AUC) technique to understand the shape of sedimentation velocity reaction boundaries that involve reversible monomer-Nmer interactions. Simulating monomer-dimer through monomer-hexamer systems as a function of concentration about the equilibrium constant allows a visual means to differentiate reaction stoichiometry by determining end points and inflexion positions. Including intermediates (eg A_1_-A_2_-A_3_-A_4_-A_5_-A_6_) in the simulations reveals the smoothing of the reaction boundary and the removal of sharp inflexions between monomers and polymers. The addition of cooperativity restores sharp boundaries or peaks to the observation and allows more discrimination in the selection of possible fitting models. Thermodynamic nonideality adds additional features when applied across wide ranges of concentration that might be appropriate for high concentration therapeutic monoclonal antibody (mAb) solutions. This presentation serves as a tutorial for using modern AUC analysis software like SEDANAL for selecting potential fitting models.

## Introduction

Beginning in the 1950’s through the 1970’s the AUC field made remarkable advances in simulating the behavior of self-and hetero-associating systems. Work from Gilbert and Gilbert (Gilbert, 1955, 1959a,b, 1960, 1963; Gilbert, Gilbert, 1973, 1978), Yphantis and Weiss (Dishon, et al., 1967, 1969; Correia, et al., 1976;), Cann and Kegeles (Bethune, Kegeles, 1961a,b; Cann, 1970, 1985; Harrington, Kegeles, 1973; Cann, Kegeles, 1974), and David Cox (Cox, 1965a,b, 1967, 1969, 1971, 1978) provided insight into the analysis of interacting systems. In 1976, the Claverie (1976) numerical solution to the Lamm equation was proposed as a visual modeling tool, to compare simulated data to AUC sedimentation velocity (SV) experimental data. Then in 1981, Todd and Haschemeyer (1981) developed the first computational method to fit AUC SV data in a rigorous manner. Subsequently, with the development of fast desk-top computers, analysis of sedimentation velocity data by numerical solutions to the Lamm equation has become a routine method for data analysis (Stafford and Sherwood, 2004). Simulations remain a powerful guide for modern investigators, in a software-dependent manner.

Many AUC studies have investigated the concentration-dependent association of therapeutic monoclonal antibodies as a potential means of selecting well behaved drugs and understanding their pharmacological properties (Hopkins, 2018, 2021; Chaturvedi, et al. 2018, 2019a,b; Liu, et al. 1995; Wright, et al. 2018a,b). Our interest is to develop AUC methods of fitting SV data to nonideal associating models. This is partially why SEDANAL was developed and tested over a wide range of protein concentrations and optical systems (Wright, et al., 2018ab, Yang, et al., 2018, Correia et al., 2020). Our goal here was to simulate and compare concerted vs sequential assembly of mAb oligomers up to hexamers. As a test system we present simulations of the Gilbert theory for self-associating monomer-Nmer mAb systems for Nmers that vary from dimer through hexamer. We then add intermediates to the simulations, for example for monomer-hexamer we simulate an A_1_-A_2_-A_3_-A_4_-A_5_-A_6_ system, keeping the association constants the same for each additional reaction step to visualize the impact of intermediates on reaction boundary shape. Nonideality and cooperativity are added to allow insight into their impact on boundary shapes. The approach is easy to implement because SEDANAL has a ModelEditor that allows complex nonideal associating models to be built, as well as a Wide Distribution Analysis WDA module that performs rapid sedimentation coefficient distribution analysis of broad SV reaction boundary shapes. This approach is easily incorporated in AUC workshops that teach the use of SEDANAL for characterization of interacting systems.

## METHODS

### Using SEDANAL to Simulate Sedimentation Velocity Data

All sedimentation velocity simulations were performed with the SEDANAL (Stafford and Sherwood, 2004; version 7.67) software package at 30,000 rpm and 100 s intervals with no added noise and enough scans to completely pellet the boundary. The position of the meniscus r_m_ was set to 5.9 cm and the cell base r_b_ set to 7.2 cm. The self-associating models for generating sedimentation velocity data were all constructed using the built-in ModelEditor (see below for the description of models; Figure S1). Simulated SV scans assuming weight extinction coefficients of 1.0 ml/mg/cm were analyzed using the WDA (Stafford, Braswell, 2004) routine in SEDANAL to generate “model-independent” sedimentation coefficient distribution functions s*g(s*) vs log(s*) for each simulated run. All WDA plots were normalized for the area under the curve and plotted as s*g(s*) vs log(s*). The molecular weight (MW) for each multimer was calculated by multiplying by the typical monomeric unit MW of 147 kDa such that dimer(A_2_), trimer (A_3_), tetramer (A_4_), pentamer (A_5_), and hexamer (A_6_) have MW’s of 284, 441, 588, 735, and 882kDa, respectively. Calculated sedimentation coefficient s_1_ values for the monomeric unit was set to be 6.50 S and the s_n_ of each subsequent multimer was calculated according to Equation 1.

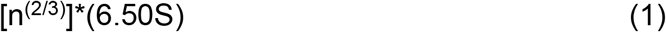

where n is the number of monomeric units. Thus, dimer (A_2_), trimer (A_3_), tetramer (A_4_), pentamer (A_5_), and hexamer (A_6_) have s_n_ values of 10.32, 13.52, 16.38, 19.01, 21.46 S, respectively. In practical terms, these can be adjusted if experimental data for oligomers or aggregates exist (Philo, 2003), or with bead modeling for hypothetical oligomer structures (see Discussion; Liu, et al. 1995; Fleming et al., 2022).

## Models

Gilbert type self-association models are monomer – Nmer reactions

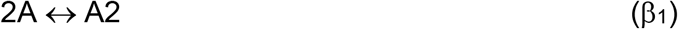

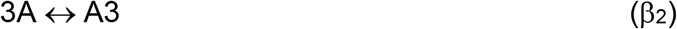

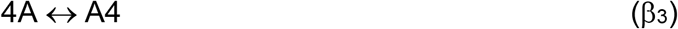

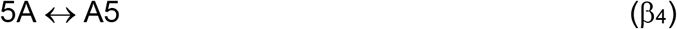

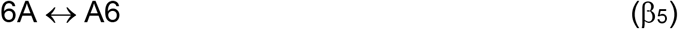

where the overall formation constants are β_1_=K_1_=K_A_, β_2_=K_1_^2^, β_3_=K_1_^3^, β_4_=K_1_^4^, β_5_=K_1_^5^ or β_1_=2.0×10^5^, β_2_=4.0×10^10^, β_3_=8.0×10^15^, β_4_=1.6.0×10^21^, β5=3.2×10^26^, respectively. This means all schemes have the same midpoint of 5 μM. For stepwise models involving intermediates

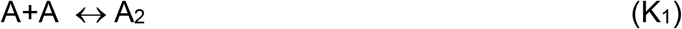

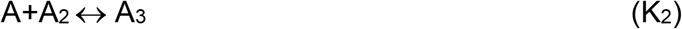

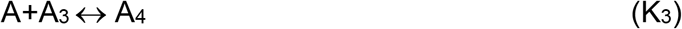

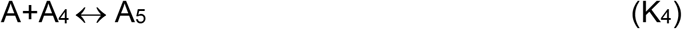

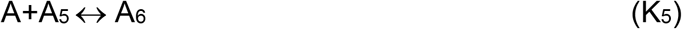

and where sequential addition of monomers assumes equal free energy; K_1_ = K_2_ = K_3_ = K_4_ = K_5_.

Nonideality is introduced as described (Stafford and Sherwood 2004; Correia et al. 2020; Correia and Stafford 2015) where k_s_ is hydrodynamic nonideality and BM_1_ is thermodynamic nonideality.

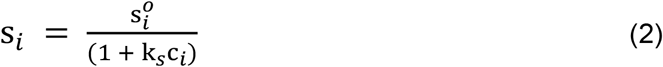

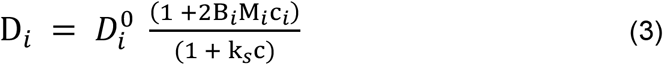

The k_s_ term can be estimated and visualized by plotting 1/s_w_ vs c according to the inverse of eq (1)

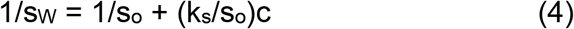

where the intercept is 1/s_o_ and the slope is k_s_/s_o_. Cooperativity is modeled by increasing K_5_ for hexamer formation by a 50 to 500-fold factor that reflects increased affinity of the final association step, and might for example reflect ring closure or other allosteric interactions (see Discussion). SEDANAL packages for data simulation (Figure S1) or analysis are available upon request.

## Results

Figure 1 is a comparison of the normalized s*g(s*) distributions for the self-association of monomeric monoclonal antibodies (mAb) into higher order Nmers, N=2 to 6, as a function of concentration. Each monomer-Nmer simulation was performed over a 100-fold range of total monomer concentrations [A] (gray curves; c_mid_/10 to c_mid_*10) and thus the peaks of each distribution range from s_1_ at low concentration to s_N_ at high concentration, where s_N_ = (n)^2/3^*s_1_. In each simulation the value of the equilibrium constant governing the single-step reversible equilibrium was taken to be the (n-1)^th^ power of 2.0×10^5^ M^-1^ or (2.0×10^5^)^(n-1)^ M^-(n-1)^. This places the midpoint of each reaction K_D_ at 5 μM and allows direct comparison between different stoichiometries (dimers to hexamers) at equal concentrations (Fig 1F). The purpose of each family of WDA plots is to visualize the shape of the reaction boundary as a function of total protein concentration. The original Gilbert theory for monomer-Nmer transport demonstrated baseline resolution of monomer peaks from polymer peaks for D=0 and N>2 (Gilbert, 1963). At concentrations of total monomer being equal to the reciprocal of the equilibrium constant (K_D_ = 5 μM) the monomer and Nmer species are clearly present as bimodal distribution curves (Fig 1F at 5 μM), where the dimer curve is skewed, the trimer curve has a pronounced inflexion near the monomer position, and for tetramer and above peaks resolve into a monomer peak centered at s_1_ and a polymer peak with an increasing degree of saturation. None of these curves are baseline resolved, although the stoichiometry of the reaction is evident in the shift of the fast boundary towards s_N_. Thus, a full Claverie simulation as implemented in SEDANAL with diffusion of a monomer-Nmer SV experiment reveals a bimodal distribution with a monomer peak (N>3) poorly resolved from a Nmer peak that approaches s_N_ as concentration approaches 10*K_D_. Approaching saturation at 10*K_D_ reflects the concerted nature of the reactions. These simulations are often described as incompatible with N monomers coming together in one kinetic step; but they are consistent with cooperative ring closure of trimers, pentamers, hexamers, or square planar or tetrahedral tetramers.

**Figure 1.**
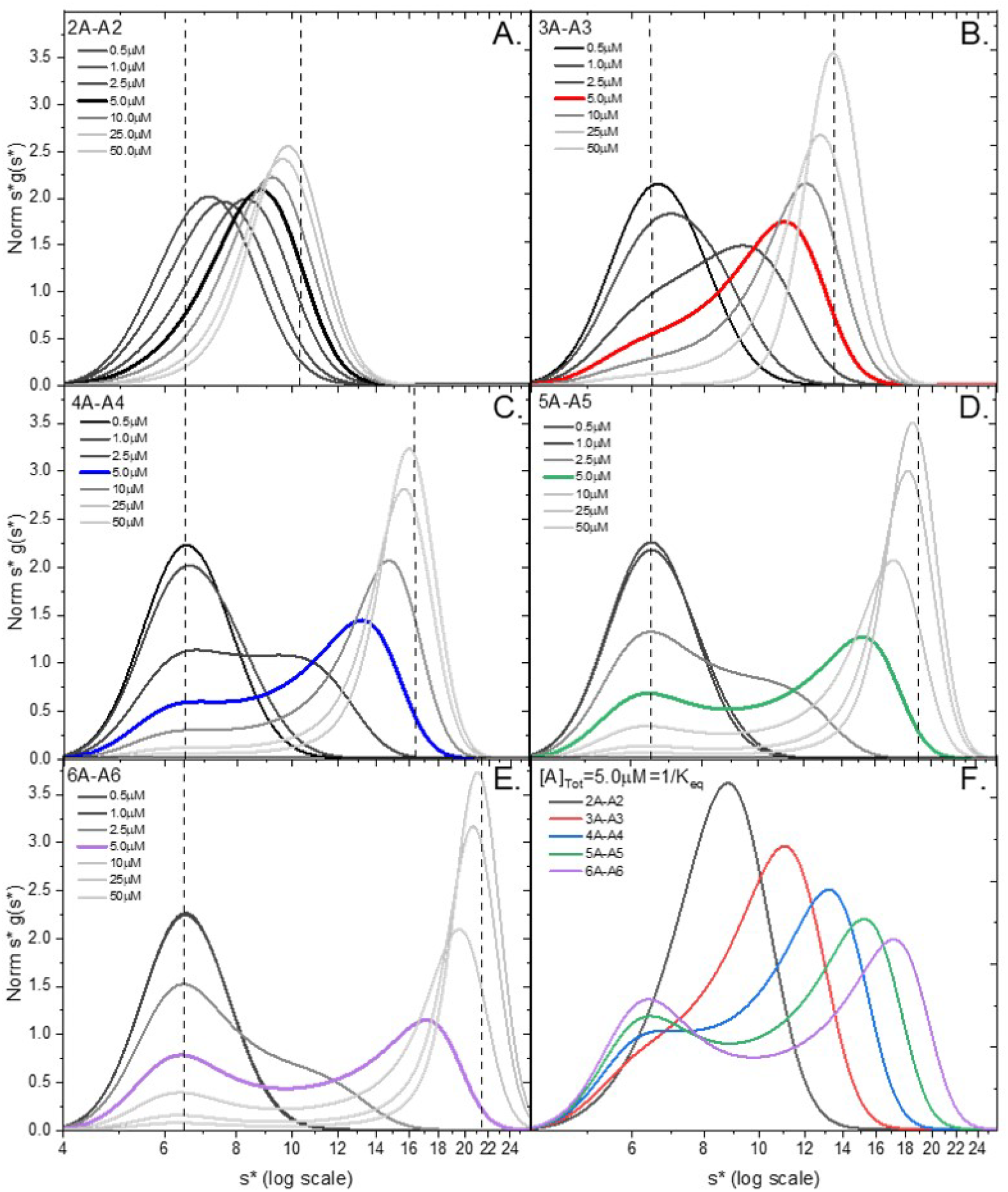
Comparison of normalized s*g(s*) sedimentation distribution analysis of concerted self-associations of monomer (nA) to multimer (An) reactions at increasing concentrations (black to gray scale). Multimers examined included dimer (Panel A, black), trimer (Panel B, red), tetramer (Panel C, blue), pentamer (Panel D, green), and hexamer (Panel E, purple). The s*g(s*) designates calculated sedimentation coefficient s values (dashed lines) for each species included in the simulation. Panel F is the comparison of the mid-point WDA plots obtained for each model at [monomer]=1/K_eq_=K_D_= 5μM.

Figure 2 presents a comparison of normalized s*g(s*) WDA plots for the ideal sequential, step-wise association of monomer (nA) into higher-order Nmers (i.e., N=2 to 6) where each step has the same value of K_A_ (i.e., K_1_ to K_5_) of 2.0×10^5^ M^-1^. Dashed lines in each panel indicate the simulated values of s_1_ for the monomer and the highest order s_N_ for the Nmer. In contrast to Figure 1, there are no inflexions and no separation into monomer and Nmer regions or peaks. Rather the s*g(s*) WDA curves shift as a concentration-dependent skewed distribution. Thus, simulated sequential models with equivalent steps do not reveal demixing (Kingsbury and Laue, 2011; Correia et al. 2020) or separation of the various oligomers. In addition, due to the need to populate intermediate sized polymers, the largest peak position is far removed from the limiting value of s_N_ at 10*K_D_. A visual comparison of Figures 1 and 2 suggestions the presence of intermediates smooths out the reaction transition and makes it more difficult to populate the largest species, the Nmer. A comparison of data at the K_D_ shows a smooth family of curves that transition towards larger s_N_ values with increasing N (Fig 2F). Thus, sequential, noncooperative models are distinguished from concerted models by the absence of bimodality. Bimodality is best observed at or near K_D_ and thus a wide concentration range is required to make these observations for experimental data.

**Figure 2.**
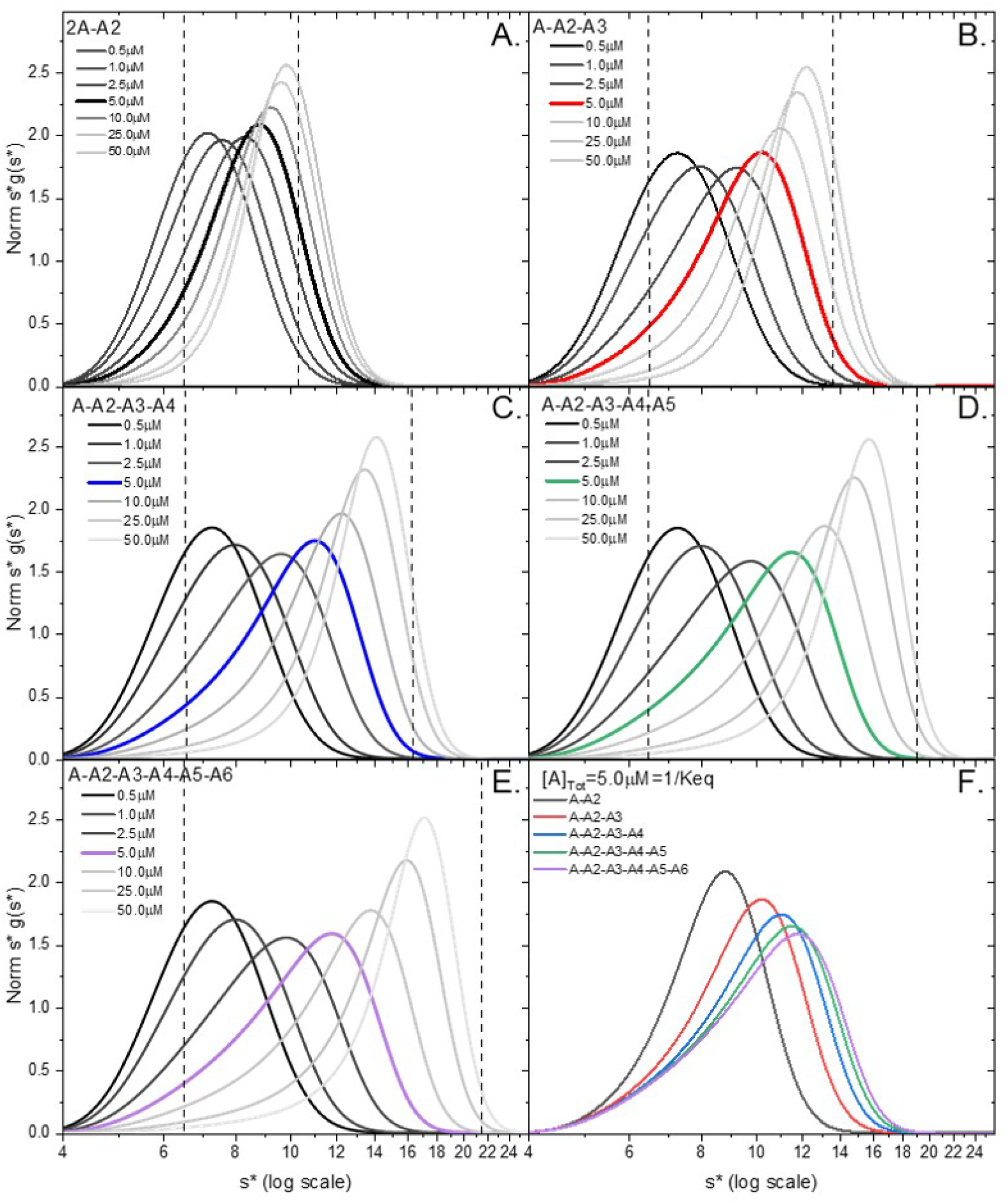
Comparison of normalized s*g(s*) sedimentation distribution (WDA) analysis of sequential multi-step self-association of monomer (A) to multimer (A2, A3, A4, A5, A6) reactions including intermediates. Multimers examined included dimer (Panel A), black), trimer (Panel B, red), tetramer (Panel C, blue), pentamer (Panel D, green), and hexamer (Panel E, purple). Panel F is the comparison of the mid-point WDA plots obtained for each model at [monomer]=1/K_eq_=K_D_=5 μM.

Therapeutic monoclonal antibodies are formulated at high concentrations, often 50 to 150 mg/ml (350 to 1000 μM). At these concentrations nonideality has a significant impact (eq 1 & 2). To demonstrate this, Figure 3 shows the effect of including nonideality in the simulations for the monomer to hexamer (A_1_ to A_6_) case for both the concerted, ie. single step (Figure 3A) and sequential models (Figure 3B). All parameters for the simulations presented in Figure 3 are identical to those presented in Figures 1 and 2 plus the inclusion of hydrodynamic k_s_ and thermodynamic BM_1_ nonideality, where k_s_ = BM_1_ = 0.01 mL/mg (10 ml/g). These are typical experimental values. To more clearly observe the impact of nonideality, the maximum concentration is increased by a factor 10, and thus the total data span is 1000-fold. In the monomer-hexamer case (Figure 3A) the observed value for S_w_ approaches s_N_ indicating saturation. In the sequential model, the system never saturates before nonideality causes a decrease in S_w_. The new feature in each of the simulations in Figure 3 is backward movement or slowing down of the higher concentrations s*g(s*) distributions (gray lines) starting at concentrations between 50-100 μM or 7-14 mg/ml. This decrease in S_w_ is due to the back flow caused by the effective volume of solvent (k_s_) displaced during sedimentation (eq 2). Thus, nonideality becomes significant at approximately 10 mg/ml. It is apparent that data collected at 25-500 μM is dominated by nonideality and if analyzed alone, would not reveal much information about the reaction mechanism without including the lower concentration data for comparison and global fitting (Correia, et al. 2020).

**Figure 3.**
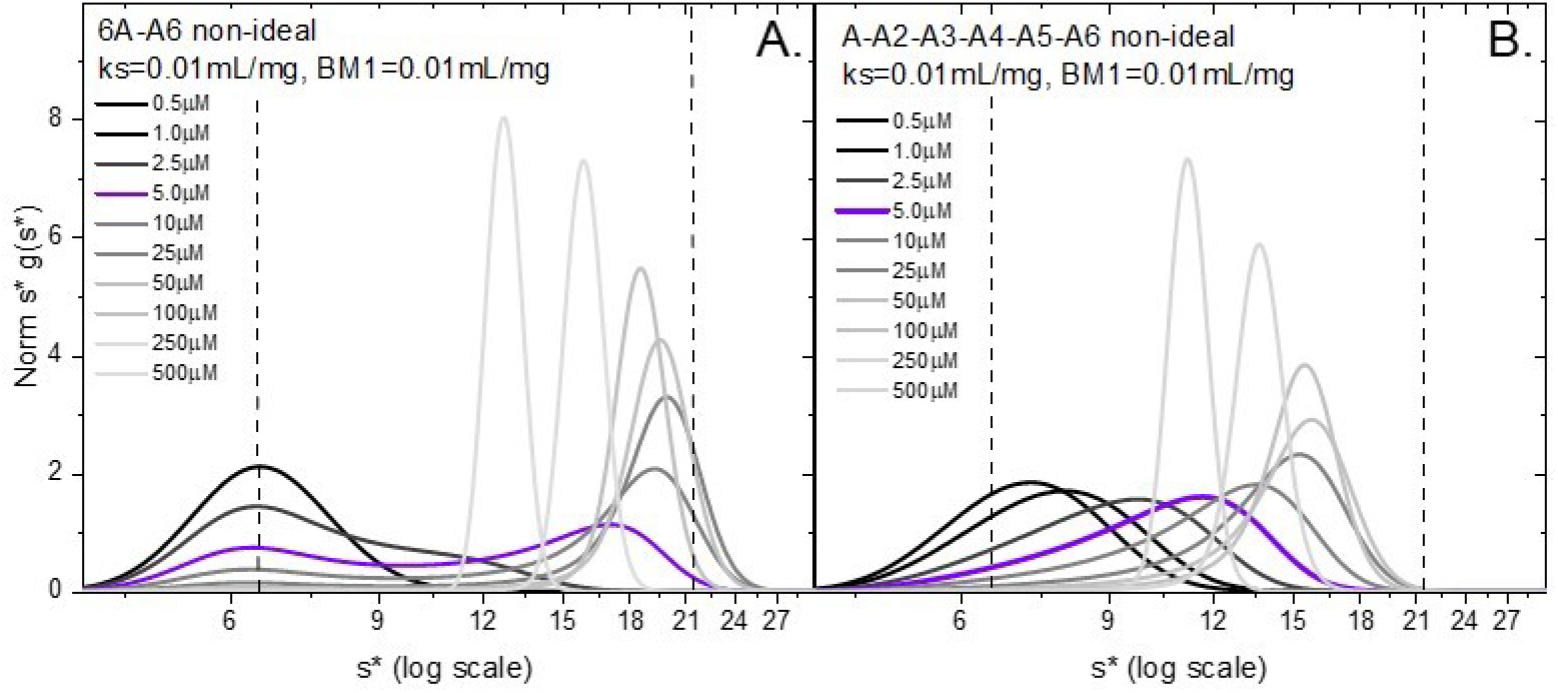
Effect of including non-ideality on the comparison of normalized s*g(s*) sedimentation coefficient distribution analysis of monomer (6A) to hexamer (A6) self-association at increasing concentrations (black to gray scale) for a concerted reaction (Panel A) and multi-step sequential reaction including intermediates (Panel B). The s*g(s*) designates calculated sedimentation coefficient distribution values (dashed lines) for beginning and end-point species included in the simulation.

To simulate cooperative binding in the sequential hexamer system, the equilibrium constant of the last reaction step (K_5_) was set to values of 10 to 500 times greater than each of the other preceding steps (K_1_-K_4_) (Figure 4A-E). In contrast to that observed in Figures 2 and 3 the resulting WDA plots of the cooperative sequential model are more clearly resolved into monomer and hexamer especially at higher levels of cooperativity (500x), but the separations are still not baseline resolved. This is especially evident in the plots at the midpoint (Figure 4F). Thus, the major distinction between concerted (Fig 1) or cooperative sequential models (Fig 4) and noncooperative sequential models (Fig 2) is the presence of a bimodal distribution. Performing simulations or experiments over a wide concentration range that includes the K_D_ is critical to seeing the evolution of this reaction boundary shape while assigning the best model.

**Figure 4.**
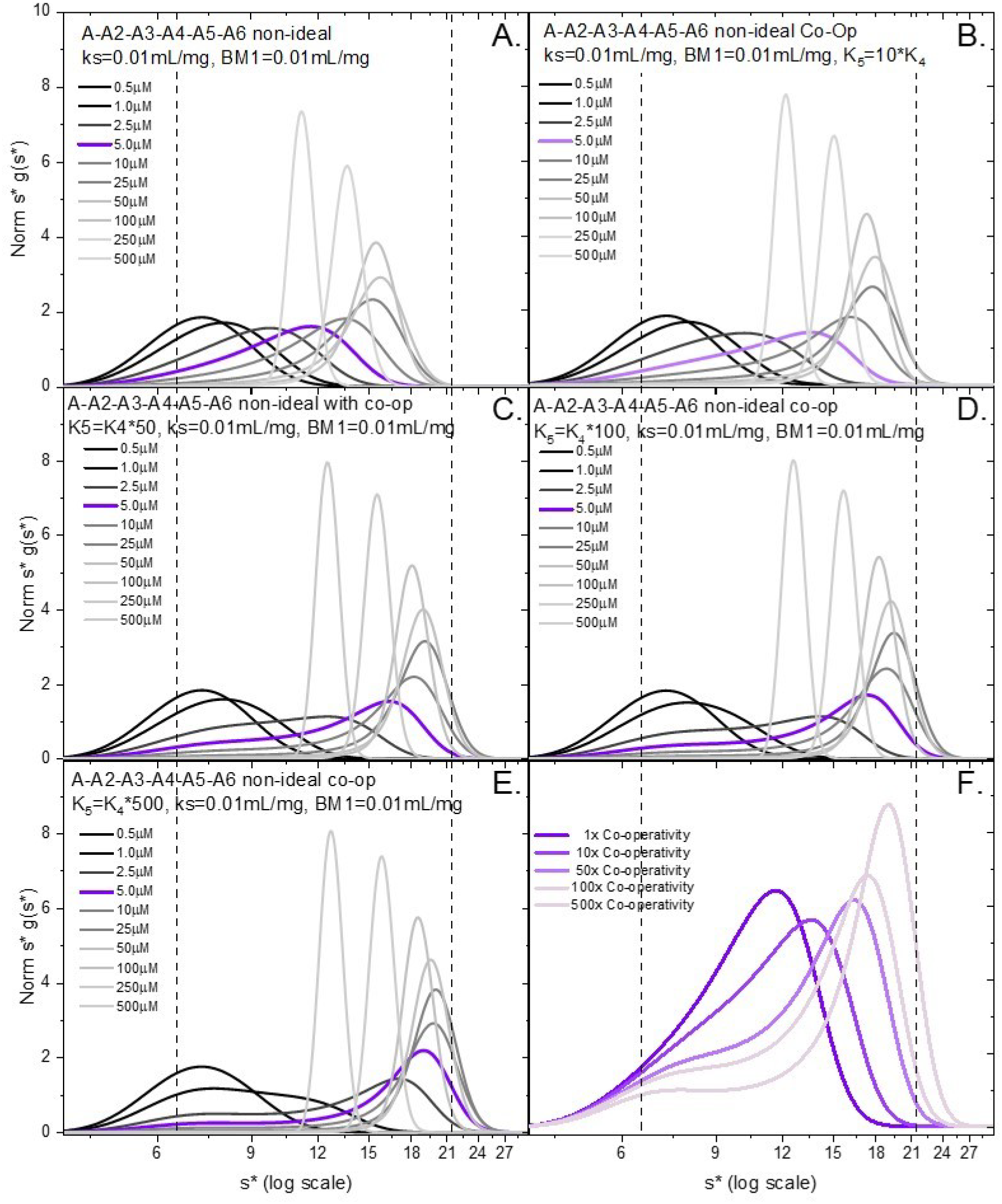
Effect of including cooperativity on the comparison of normalized s*g(s*) sedimentation distribution analysis of nonideal sequential monomer (A) to hexamer (A6) by self-association with intermediates at increasing concentrations (black to gray scale). The s*g(s*) designates calculated sedimentation coefficient distribution values (dashed lines) for monomeric and hexameric end-point species included in the simulation.

Figure 5B summarizes the resulting weight average S_w_ values of the monomer hexamer systems extracted from the normalized model-independent WDA plots as a function of initial concentration (c_o_) for the fully cooperative and nonideal stepwise formation of hexamer. Data are cast as S_w_ vs logc_o_ (mg/ml) to observe the presence of cooperativity, which exhibits a steeper rise in S_w_ relative to the sequential model. The nonideal data have a hint of curvature or overshoot in the plots. This can in principle be more clearly seen in the 1/S_w_ vs c plot (Figure 5D) where the data curve upward and become linear with a slope of (k_s_/s_o_) (eq 4). This suggests the need to include nonideality, and the ability to analyze nonideality is dictated by the K_D_ and the span of the data. However, in this case the data do not span a wide enough region to see the impact of nonideality on a 1/S_w_ vs c plot. To demonstrate this, similar data were simulated at weaker and tighter affinity, 50 μM (Figure 5A,B) and 0.5 μM (Figure 5E,F). The weaker affinity data demonstrate significant over shoot caused by the nonideality above the midpoint of 7.3 to 73 mg/ml (Figure 5A); the tighter affinity data corresponding to K_D_ = 0.5 μM are free of nonideality effects because the data never exceed 0.73 mg/ml (Figure 5E). These plots are reproduced as Fig S2 over a much broader span of concentration to demonstrate that nonideality becomes significant over the same maximum concentration range. This is more evident in panels S2B, D, and F where the linear portion of the curve becomes well-determined. These results are limited to 1000-fold and 10000-fold simulations and are unlikely to be explored experimentally. Nonetheless it gives a user an understanding of concentration effects and the importance of simulating and collecting experimental data over the widest range possible.

**Figure 5.**
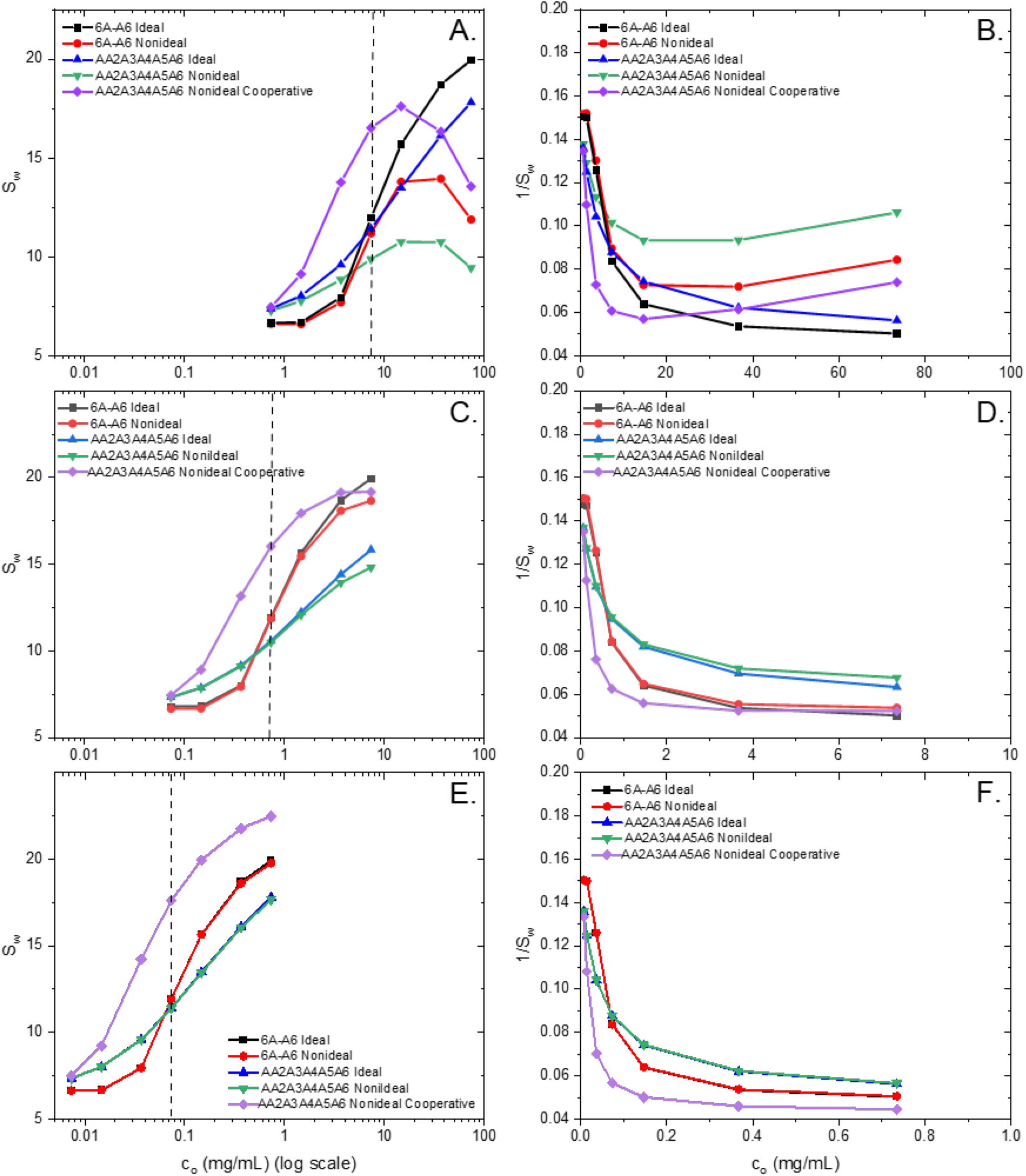
Summary of the weight averaged S_w_ values for concerted and sequential monomer hexamer simulations versus initial concentration log(c_0_) in mg/mL centered around K_D_=1/K_A_. Panels A, C, and E panels plot the S_w_ versus logc_o_ for K_eq_=2.0×10^4^ (Panel A), K_eq_=2.0×10^5^ (Panel C), and K_eq_=2.0×10^6^. Panels B, D, and F plot the reciprocal of S_w_ (i.e., 1/S_w_) versus c_o_ for K_eq_=2.0×10^4^ (panel B), K_eq_=2.0×10^5^ (panel D), and K_eq_=2.0×10^6^ (panel F). The dashed vertical line in the panels A, C, and E mark the center concentration=K_D_. The purple lines includes 500x cooperativity in K_5_ and simulations including non-ideality (red and green/purple) set ks and BM1 to 10 mL/mg each.

## Discussion

Our goal here is to create a general path for simulating SV reaction boundaries for self-associating mAbs. The approach is based upon the 1950’s and 60’s Gilbert Theory for self-association but utilizes the full numerical solution to the Lamm equation (Claverie, 1976) as implemented in SEDANAL, including diffusion, nonideality and cooperativity. Graphical representations of the reaction boundary shapes of different monomer-Nmer models are presented as families of wide distribution or WDA plots as a function of concentration. This approach of using WDA to determine the sedimentation coefficient distribution is quick and preferable to DCDT because it uses all the data scans and accurately captures a wide sedimentation distribution spanning monomers to hexamers; DCDT works best with a narrow range of scans and thus cannot accurately capture reversible monomer-hexamer boundaries without significant broadening. Sedfit or c(s) analysis is inappropriate because it is not designed to represent reversible interacting systems, and it tends to produce peaky graphics that do not reveal an accurate shape of a reversibly interacting boundary (Schuck, 2005). This can be seen in a direct comparison of WDA, DCDT+ and c(s) analysis of the hexamer data at 5 μM (Figure S6A, C). To further emphasize this the concentration of each species is plotted vs radial position to capture the composition of reacting boundary early in the simulation (scan 75/400; Figure S6B,D). The c(s) peaks clearly do not correspond to actual boundaries in the concentration plots. It should be emphasized that every radial position in the concentration vs radius plots (Figure S6B,D) obeys the mass action equilibrium simulated (Figure S1).

This approach of simulating and presenting gradients or boundary shapes, was reemphasized by David Cox for nonideal trimer systems in 1971, pointing out the presence of minima and inflexions as criteria for evaluating the best models, while still being approximate asymptotic solutions to the Lamm equation. Claverie’s numerical approach captured the full solution to simultaneous sedimentation and diffusion, but was also meant to be a graphical approach for comparison to experimental data. Todd and Haschemeyer (1981) offered the solution to the inverse problem by comparing data and Claverie simulations as a least squares best fit. The computational capacity to globally fit real data had to wait until the development of a platform that integrates multiple concentrations, multiple signals, and complex models that involve nonideal, self-and hetero-association (Stafford and Sherman, 2004). And yet the selection of best models is still a challenge for new users who lack experience with numerical methods and an understanding of reversible associating systems. A need for a wide range of concentrations or a wide range of ratios for hetero-systems (to be discussed elsewhere) is not fully appreciated. Sufficiently high stock concentrations of mAb are often the constraining experimental limitation. Here we show that in silico solutions are unlimited in scope and allow a vision of the full evolution of reaction dynamics. To demonstrate this here we present a range of concentrations and models that are appropriate for reacting mAb systems.

Many AUC studies have investigated the concentration-dependent association of therapeutic antibodies as a potential means of selecting well behaved drugs and understanding their pharmacological properties. Our interest is to develop AUC methods of fitting SV data to nonideal associating models. This is partially why SEDANAL was developed and tested over a wide range of concentrations and optical systems (Correia and Stafford, 2009; Correia, et al. 2009; Wright, et al., 2018ab, Yang, et al., 2018, Correia et al., 2020). Our goal here was to simulate and compare cooperative vs sequential assembly of mAb oligomers up to hexamers. Many of the published studies struggle with fitting to realistic models, and report a wide range of best fitted models, including weak dimerization, dimer-tetramer-hexamer, and isodesmic (Hopkins, et al., 2018, 2021; Wright et al. 2018a,b). We propose the ability to simulate an equally wide range of models to explore distinct features of reaction boundaries makes this process more productive and accurate. For example, our results suggest bimodal boundaries with inflexions and partially resolved peaks are suggestive of cooperative models. Thus, a dimer-tetramer-hexamer model that displayed bimodality will require a cooperative best fit, where for eg. K_2_ < K_4_ < K_6_ to be consistent with the experimental data. The exact cooperative model will depend upon the nature of the interactions (see Liu et al. 1995 for example). Isodesmic models are essentially sequential models with equal affinity and thus, consistent with our simulations, show skewed reaction boundaries without any inflexions or bimodality. These features provide additional confidence in the best fitted results.

An advantage of simulations is that we are not constrained by limitations of a particular optical system. These results are best applied to interference signals that can easily span 0.01 to ∼50 mg/mL. Absorbance measurements are generally more limiting in concentration range due to the large extinction coefficient of typical mAbs at 280 nm (∼1.5 mL/mg/cm), where 0.1 mg/ml to 7 mg/ml are possible by combining 12 mm, 3 mm, or 1.5 mm cells (Nanolytics). Thus, Abs would only allow sampling the lower half of the reactions in Figs 1 and 2. Collecting data near the absorbance minima at 255 nm expands this range typically by a factor of 2 or more. FDS optics offers the potential for data collection from low nM (.001 mg/ml) to 100-150 mg/mL in tracer mode (Kingsbury, Laue 2011; Correia, et al. 2020). But the conclusions about reaction boundary shapes will still be valid. The main limiting feature in high concentration FDS data is the dominant role nonideality plays above 10-20 mg/mL. Thus, proper fitting of high concentration FDS data requires a broad range of concentrations that allows both association and nonideality properties to be extracted (Correia, et al. 2020). Nonideal data alone masks the association process. We have previously pointed out that fitted k_s_ values that are too small and inconsistent with calculated values for mAbs (11 ml/g) are suggestive of weak association and the need to add association to the best model (Yang, et al. 2018; Fleming, et al. 2022). The downward portion of the curves in Fig S2B,D,F correspond to the association phase, while the linear portion of these curves correspond to the nonideal region of the curves. Note in Figure 5B association and nonideality overlaps since the weak association will require high concentration data. In Figure 5D there is only a small region where nonideality contributes and fitting data below 5 mg/ml may not need nonideality to improve the fit. The data in Fig 5F exhibits no nonideality since it is all at very low concentrations. This is why the ability of SEDANAL to extract association properties in the presence of nonideality is so critical. Data must be fit over a wide range of concentrations to both “see” the association behavior around K_D_, while also “seeing” the nonideal behavior at high concentrations if present. This has been the focus of many of our prior publications (Correia, Stafford 2009; Correia, et al. 2009, 2020). Fitting over various spans of the data will reveal how well determined the various parameters are (Tables 1 and 2 in Correia et al. 2020).

In conclusion, SEDANAL is a powerful AUC software package to simulate assembly models that may resemble experimental data. It is critical that the experimental data span a wide concentration range about the K_D_. This allows the best opportunity to match features between simulated models and experimental data. This applies to situations where one is trying to discriminate between cooperative and noncooperative systems, especially systems with multiple species that might exhibit bimodal sedimentation coefficient distributions. This allows one to rule out models that are inappropriate and unlikely to fit the data. Globally fitting data to complex models can take extensive computational time, especially when performing Fstat or Bootstrap analysis, and the fewer models tested will speed up the best selection process. Nonideality is often required in best fits and its inclusion can be anticipated by noting the magnitude of k_s_c, which for typical mAb systems is 0.01*c, which makes 1 + 0.01*c in equation 2 equal to 1.1 at 10 mg/ml. Many systems with more exaggerated, rod like shapes can have k_s_ values > 100 ml/g (Creeth and Knight, 1965) which means nonideality becomes significant at 1 mg/ml. Comparing simulated WDA plots along with s_w_ and 1/s_w_ vs c plots allows estimates of K_D_ and k_s_ values and thus better initial guesses for nonlinear least squares. We have previously published approaches for fitting high concentration FDS data (Correia, et al., 2020) that provide insight into the impact of k_s_ and BM1 on boundary shapes. The utility of SEDANAL is the ability to extract model-dependent equilibrium constants and nonideality parameters that properly reflect attractive and repulsive interactions over a wide range in concentration.

## Acknowledgements

We thank Dave Bain, Sharon Lobert, Pat Fleming and Walter Stafford for constructive comments and suggestions. Tutorial modules for workshop presentations are available from JJC and Walter Stafford.

## Supplemental Figures

**Figure S1.**
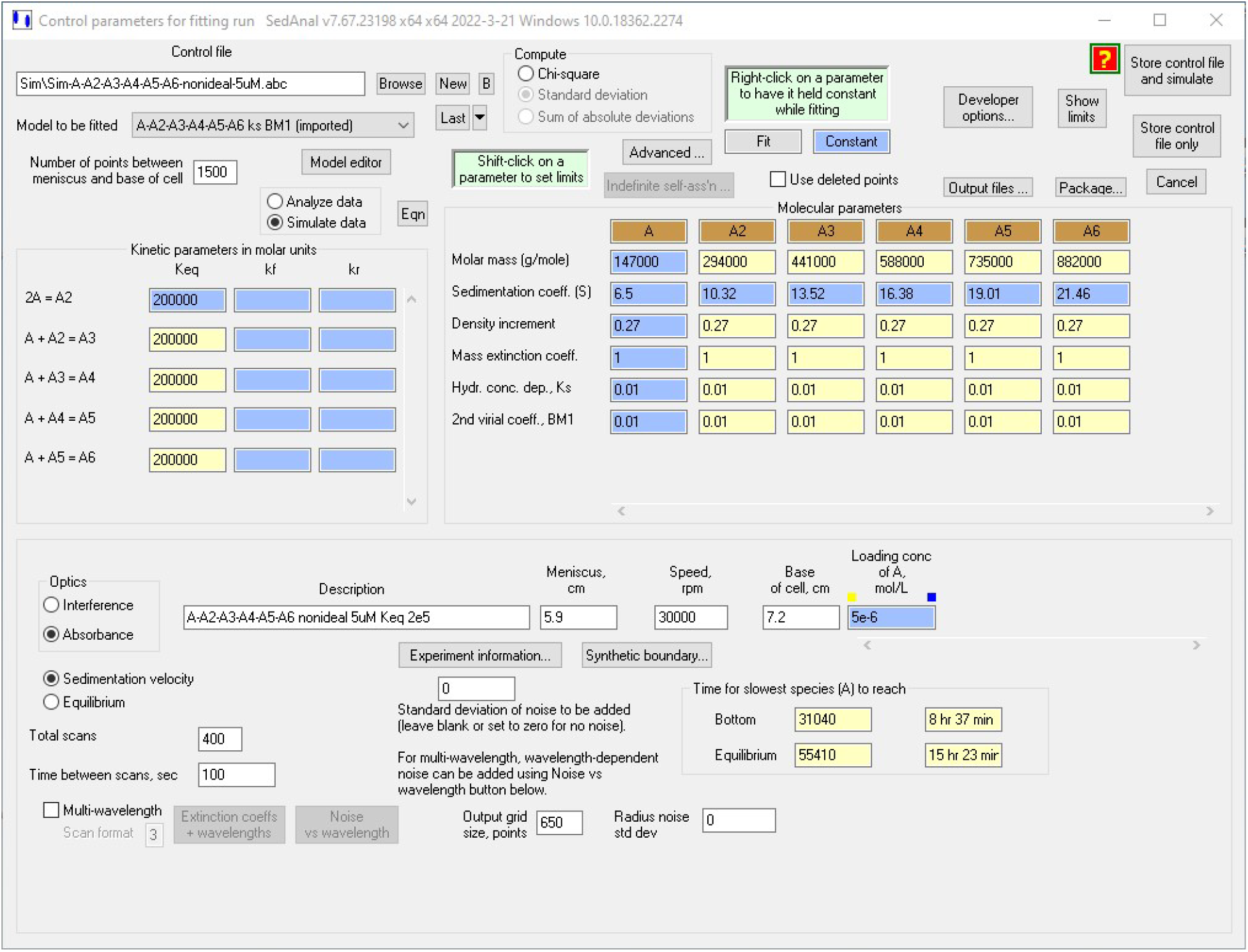
The ABC file for simulating nonideal sequential hexamer formation with each K = 2e5, ks = 10 ml/g, and BM1 = 10 ml/g at 5 uM.

**Figure S2.**
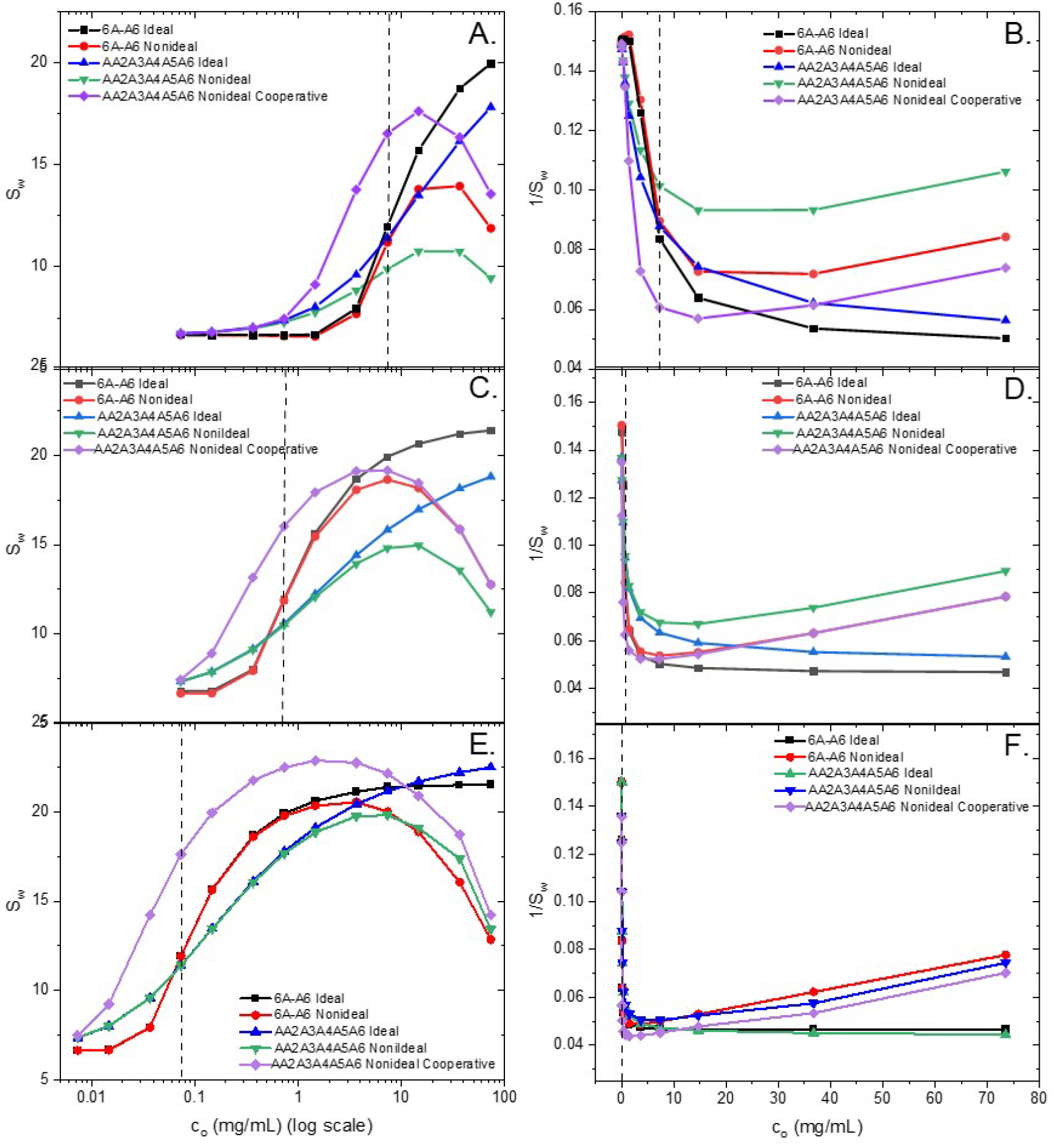
Summary of the full concentration range for the weight averaged S_w_ values versus initial concentration (c_0_) in mg/mL. Panels A, C, and E panels plot the S_w_ versus logc_o_ for K_eq_=2.0×10^4^ (Panel A), K_eq_=2.0×10^5^ (Panel C), and K_eq_=2.0×10^6^. Panels B, D, and F plot the reciprocal of S_w_ (i.e., 1/S_w_) versus c_o_ for K_eq_=2.0×10^4^ (panel B), K_eq_=2.0×10^5^ (panel D), and K_eq_=2.0×10^6^ (panel F). The dashed vertical line in panels (A, C, and E) mark the center concentration=K_D_. The purple lines is the result of including 500x cooperativity in K_5_ and simulations including non-ideality (red and green/purple) with ks and BM1 to 10 mL/mg each. The data for filling in the concentration regions is from the simulations shown in Figures S3, S4 and S5.

**Figure S3.**
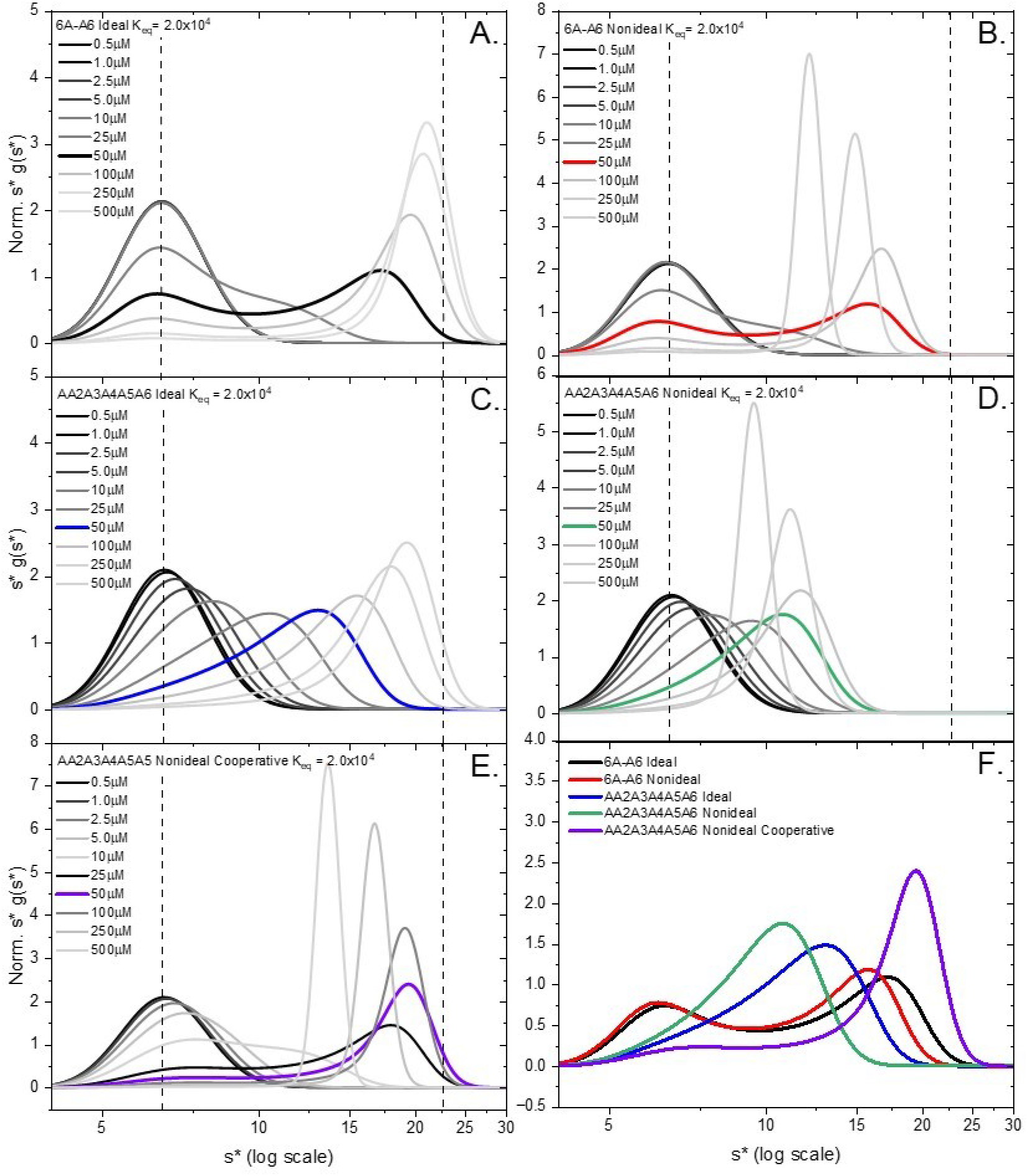
Normalized WDA plots comparison of hexameric models for K_eq_=2.0×10^4^. Concerted (6A-A6) ideal (panel A) and non-ideal (panel B) and sequential (A-A2-A3-A4-A5-A6) ideal (panel C) and non-ideal (panel D). Non-ideality was modeled with ks and BM1=10mL/mg and 500x cooperativity (panel E). Panel F plots the midpoint concentration equal to 1/K_eq_=K_D_ for each case.

**Figure S4.**
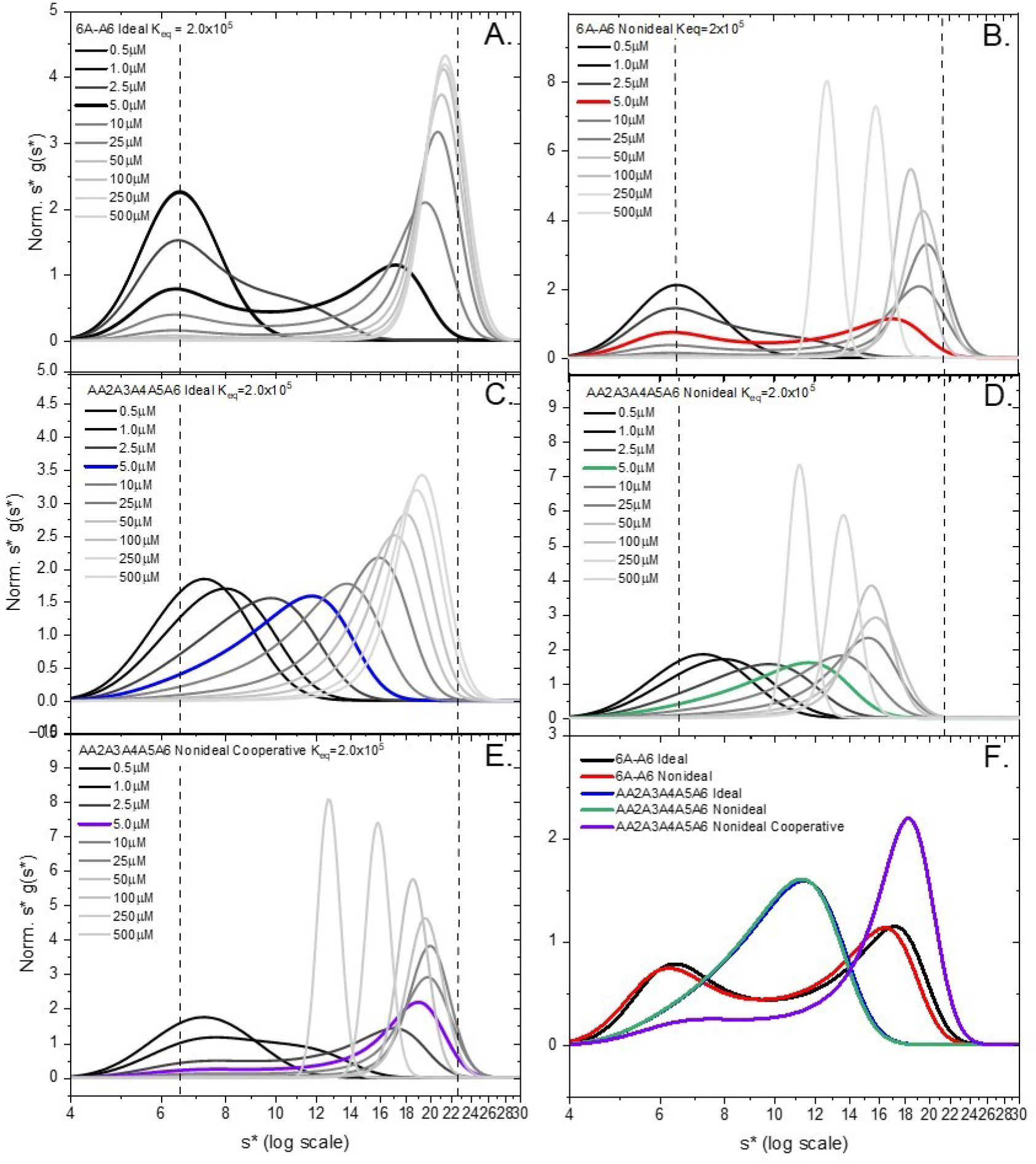
Normalized WDA plots comparison of hexameric models for K_eq_=2.0×10^5^. Concerted (6A-A6) ideal (panel A) and non-ideal (panel B) and sequential (A-A2-A3-A4-A5-A6) ideal (panel C) and non-ideal (panel D). Non-ideality was modeled with ks and BM1=10mL/mg and 500x cooperativity (panel E). Panel F plots the midpoint concentration equal to 1/K_eq_=K_D_ for each case.

**Figure S5.**
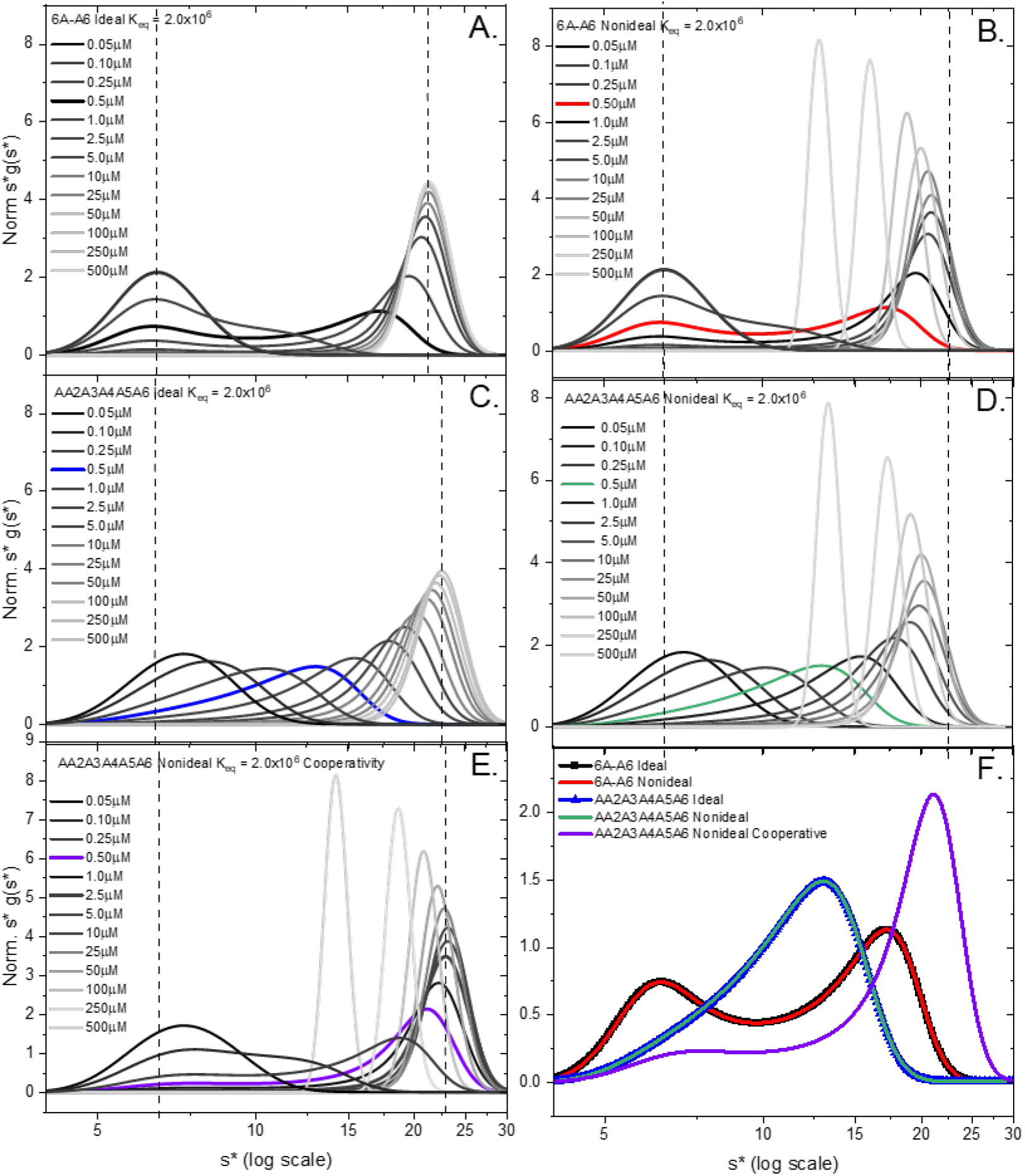
Normalized WDA plots comparison of hexameric models for K_eq_=2.0×10^6^. Concerted (6A-A6) ideal (panel A) and non-ideal (panel B) and sequential (A-A2-A3-A4-A5-A6) ideal (panel C) and non-ideal (panel D). Non-ideality was modeled with ks and BM1=10mL/mg and 500x cooperativity (panel E). Panel F plots the midpoint concentration equal to 1/K_eq_=K_D_ for each case.

**Figure S6.**
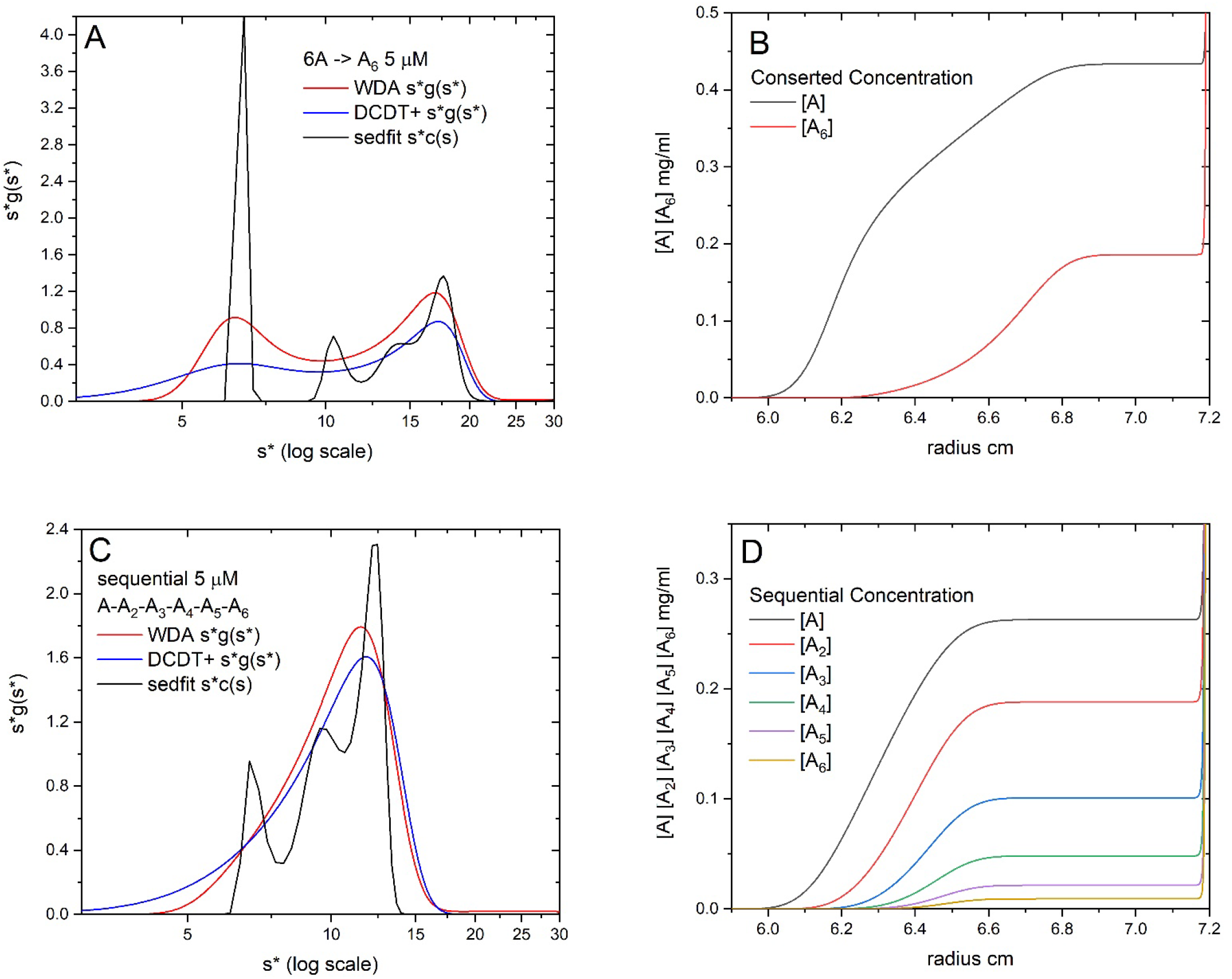
Overlay of WDA, DCDT+ and Sedfit analysis of data simulated at 5 uM for a A) concerted and C) sequential monomer hexamer system. Concentrations are plotted vs radius at some early time in the simulation for each species in the B) concerted and D) sequential model. These represent the composition of the reacting boundary where at each radial position the equilibrium is maintained throughout the simulation. The sum of the concentrations in panels B and D are described by the WDA s*g(s*) distributions (red lines) in panels A and C.

